# The unstructured linker arms of MutL enable GATC site incision beyond roadblocks during initiation of DNA mismatch repair

**DOI:** 10.1101/463133

**Authors:** Yannicka SN Mardenborough, Katerina Nitsenko, Charlie Laffeber, Camille Duboc, Enes Sahin, Audrey Quessada-Vial, Herrie HK Winterwerp, Titia K Sixma, Roland Kanaar, Peter Friedhoff, Terence R Strick, Joyce HG Lebbink

**Affiliations:** Department of Molecular Genetics, Erasmus University Medical Center, Rotterdam, the Netherlands; Institut Jacques Monod, CNRS, UMR7592, University Paris Diderot, Sorbonne Paris Cité F-75205 Paris, France; Oncode Institute, the Netherlands; Division of Biochemistry, Netherlands Cancer Institute, Amsterdam, the Netherlands; Institute for Biochemistry, Justus-Liebig University, Giessen, Germany; Ecole Normale Supérieure, Institut de Biologie de l’Ecole Normale Superieure, CNRS, INSERM, PSL Research University, 75005 Paris, France; Programme “Equipe Labellisée”, Ligue Nationale contre le Cancer; Department of Radiation Oncology, ErasmusMC, Rotterdam University Medical Center, the Netherlands

## Abstract

DNA mismatch repair (MMR) maintains genome stability through repair of DNA replication errors. In *Escherichia coli*, initiation of MMR involves recognition of the mismatch by MutS, recruitment of MutL, activation of endonuclease MutH and DNA strand incision at a hemimethylated GATC site. Here we studied the mechanism of communication that couples mismatch recognition to daughter strand incision. We investigated the effect of catalytically-deficient Cas9 as well as stalled RNA polymerase as roadblocks placed on DNA in between the mismatch and GATC site in ensemble and single molecule nanomanipulation incision assays. The MMR proteins were observed to incise GATC sites beyond a roadblock, albeit with reduced efficiency. This residual incision is completely abolished upon shortening the disordered linker regions of MutL. These results indicate that roadblock bypass can be fully attributed to the long, disordered linker regions in MutL and establish that communication during MMR initiation occurs along the DNA backbone.

## Introduction

DNA mismatch repair (MMR) is an evolutionary conserved process responsible for detection and repair of base-base mismatches and insertion-deletion loops that have escaped the proofreading capacity of DNA polymerase during DNA replication^1^. Functional loss of MMR results in a mutator phenotype and, in humans, in a predisposition to develop cancer (Lynch syndrome)^2^. The first step in DNA MMR, recognition of the replication error, is carried out by MutS (in prokaryotes ^3,4^) or one of its homologs (in eukaryotes ^5,6^). After ATP-dependent mismatch verification and its concomitant conformational change in MutS ^7-10^, MutL (or MutLα) is recruited ^10-14^. In an ATP-dependent manner this complex activates a latent endonuclease activity which either resides in an independent protein (MutH in γ-proteobacteria such as *Escherichia coli* ^15-17^ or within the C-terminal part of MutL/MutLα ^18,19^. This nuclease activity is responsible for creating nicks in the newly synthesized daughter strand, which subsequently allows removal of this strand including the incorrectly incorporated nucleotide, and correct resynthesis ^1,20^.

While the exact molecular mechanism for strand discrimination in most organisms remains to be fully determined ^21-24^, it is established that γ-proteobacteria use the transient hemi-methylated state of GATC sites immediately after replication to distinguish parental from daughter strand ^25^. GATC sites are methylated on the adenine base by the action of *dam* methylase, which lags behind the replication fork by about one minute ^26,27^. Within this limited time window, the MutH endonuclease can introduce a nick 5’ of the deoxyadenosine in the unmethylated strand of the hemimethylated GATC sites in the DNA duplex ^28^, which is the strand containing the misincorporated nucleotide.

Because the replication error and strand discrimination signal are located at different sites and in different orientations on the DNA, MMR employs a directional communication mechanism to ensure that daughter strand incision and excision results in removal of the fragment of the DNA that includes the replication error, rather than excision tracts being directed away from the mismatch ^29,30^. A functional study of note that addresses this communication ^31^ describes how the placement of a nuclease-deficient variant of EcoRI (EcoRI_E111Q_) as roadblock in between a mismatch and a GATC site on a linear DNA fragment significantly reduces strand incision, supporting models that invoke communication along the DNA backbone. Events that escape inhibition were attributed to microscopic dissociation of the roadblock from its recognition site, to formation of loops through space involving segments of the DNA on different sides of the roadblock, or both of these mechanisms. Recent observations from single molecule approaches in which *E. coli* MutL and yeast MutLα were both observed to be able to bypass other proteins present on the DNA ^32-34^, prompted us to investigate whether this behavior could underlie the ambiguity in the interpretation of the observed roadblock bypass during MMR strand incision.

Here we describe two new functional assays (one ensemble and one single molecule approach) in which GATC site incision by MMR proteins can be quantitatively correlated to the absence and presence of different roadblocks in between mismatch and GATC site. Using a MutL variant carrying shortened disordered linkers in between its globular domains, we find that a model which invokes communication along the DNA backbone is sufficient to describe MMR initiation.

## Results

### A quantitative assay to monitor MMR strand incision at individual GATC sites

To explore the mechanism by which MMR proteins communicate between mismatch and multiple hemimethylated GATC sites during initiation of DNA mismatch repair, we quantitatively investigated strand incision using linear substrates containing a single G/T mismatch (or its corresponding A/T base pair) flanked by two hemimethylated GATC sites and an Alexa^647^ fluorophore (Figure 1A). Circular substrates ^35^ were linearized using ScaI, mixed with 50 nM MutS, 50 nM MutL and 25 nM MutH in the presence of ATP, and formation of reaction products carrying the fluorophore was analyzed using denaturing gel electrophoresis, which allowed identification of nicking events at individual GATC sites. Efficient incision of both GATC sites was observed on GT#2, carrying a G/T mismatch, while activity on homoduplex AT#2 substrate was negligible under these conditions (Figure 1B).

**Figure 1:**
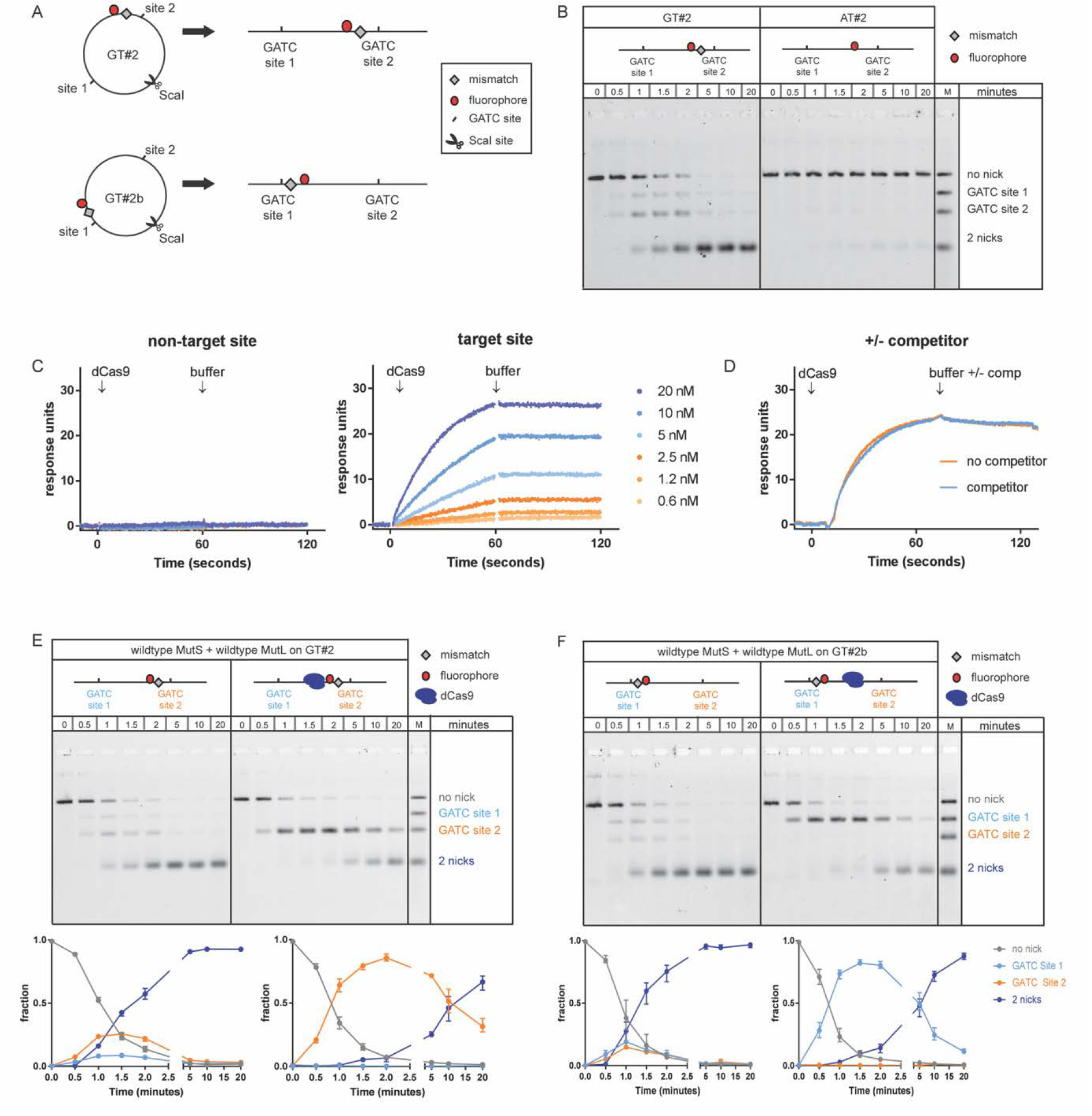
Effect of dCas9 roadblock on incision of GATC sites during initiation of DNA mismatch repair. A) DNA substrates contain a single G/T mismatch and Alexa^647^ fluorophore at different positions flanked by two hemi-methylated GATC sites and are linearized using ScaI. B) Time courses for strand incision by 50 nM MutS, 50 nM MutL and 25 nM MutH on 0.5 nM GT#2 (left panel; substrate carrying mismatch) or AT#2 (right panel; control containing the corresponding A/T Watson-Crick base pair). Reaction products are separated using gel electrophoresis under denaturing conditions and visualized using Alexa^647^ fluorophore emission. C) Surface plasmon resonance of dCas9-RNA complex binding to target and non-target sites using Biacore T100. Left panel: dCas9-RNA_A complex (0.6-20 nM) injected over 60-bp duplex DNA carrying target site C. Right panel: dCas9-RNA_C complex (0.6-20 nM) injected over 60-bp duplex DNA carrying site C. D) Control addressing putative rebinding of dCas9-RNP to target site during dissociation. dCas9-RNA_C (20 nM) was injected over the surface, and dissociation was monitored in the absence (red) or presence (green) of excess non-biotinylated duplex containing target site C (competitor). E, F) Time courses for hemimethylated GATC site incision on 0.5 nM GT#2 (panel E) and 0.5 nM GT#2b (panel F) by 50 nM MutS, 50 nM MutL and 25 nM MutH in the absence (left) and presence (right) of a dCas9 roadblock placed on target site C in between the GATC sites. Reaction products are separated using gel electrophoresis under denaturing conditions and visualized using the Alexa^647^ fluorophore. Graphs underneath the gels show fractions of product containing no nick (gray), a nick at GATC site 1 (light blue), a nick at GATC site 2 (orange) and 2 nicks (dark blue). Data points with error bars represent the mean values and range of three independent experiments.

On substrate GT#2, we observed formation of small amounts of reaction intermediates containing a single incision at either GATC site 1 or site 2, and progressive accumulation of reaction product in which both GATC sites were incised (Figure 1B and 1E; left panel). This rapid conversion of single-nicked intermediates into double-nicked products on linearized substrate is similar to what we have observed previously on circular GT#2 ^35^. However, while on circular GT#2 GATC site preference was undetectable (Supplemental Figure S1), on linear GT#2 we observed a preference for nicking GATC site 2, which is close to the mismatch (Figure 1E left panel; Supplemental Figure 1). Upon changing the location of the mismatch to a position that is now closer to GATC site 1 in an otherwise identical configuration (substrate GT#2b in Figure 1A), this site preference is reversed (Figure 1F left panel; Supplemental Figure S1). These observations indicate that the presence of DNA ends, the distance between mismatch and GATC site, and possibly the distance of these elements from DNA ends, differentially influence the efficiency by which individual GATC sites become incised.

### Slow bypass of dCas9 roadblock during MMR initiation

Modrich and colleagues have reported reduced strand incision efficiencies upon placement of a nuclease-deficient variant of EcoRI (EcoRI_E111Q_) as roadblock in between mismatch and GATC site on a linear DNA substrate ^31^. However, significant remaining incision activity beyond this roadblock was observed, attributed at least in part to the EcoRI_E111Q_ roadblock vacating its target site ^31^. To address whether incomplete block formation is indeed the reason for the observed residual incision activity, we investigated whether the Cas9 ribonucleotide-protein (RNP) complex from *Streptococcus pyogenes* ^36^ could be used as a more stable site-specific roadblock. Cas9 forms an RNP complex with tracrRNA and a specific crRNA which it uses to form an R-loop with a complementary DNA target site sequence. R-loop formation relies on the presence of a PAM (5’-NGG) sequence on the non-target strand immediately upstream of the recognition sequence. For our purpose, we used a catalytically dead mutant (dCas9) containing single point mutations (D10A and H840A) in both nuclease domains. The Cas9 RNP complex forms a larger roadblock than EcoRI_E111Q_ (~195 kDa instead of 62 or 124 kDa depending on EcoRI_E111Q_ oligomeric state) and forms sequence-specific interaction with the DNA via Watson-Crick base pair formation within the R-loop rather than direct protein-DNA interactions. RNP-DNA complex formation was probed using surface plasmon resonance (SPR) analysis on 60-bp duplex DNA carrying target site C (see Materials and Methods), and was found to occur with high stability and specificity (Figure 1C). The observed stable complex formation is not caused by rebinding of dissociated dCas9 to adjacent DNA molecules during the wash phase, as inclusion of a large excess of competitor DNA carrying the target site C sequence during the buffer wash does not have an effect on the observed dissociation rate (Figure 1D). Experiments with longer dissociation times on different target sites (not shown), as well as reported observations ^37,38^ indicate that a typical dCas9-target site complex has a half-life of at least 6 hours. This is considerably longer than the reported 40 minutes for EcoRI_E111Q_ ^31^. This combination of larger size, longer residence time and different mode of interaction prompted us to use the dCas9 RNP complex as roadblock during strand incision.

We placed the dCas9 RNP complex as a site-specific roadblock on target site C, which is located in between the two GATC sites on our MMR substrates. In the presence of roadblock C, substrate GT#2 was incised at GATC site 2 located close to the mismatch with an efficiency similar to that on unblocked DNA (Figure 1E right panel). No reaction intermediates containing an incision only at site 1 beyond the roadblock were observed. Likewise, substrate GT#2b with the mismatch close to GATC site 1, was incised at site 1 with similar efficiency as on unblocked DNA, with no reaction intermediates containing only a nick at site 2 beyond the roadblock being produced (Figure 1F right panel). However, on both substrates roadblock bypass was observed, indicated by the appearance of product with incisions at both sites, with this second nick being introduced 3-5-fold slower than on unblocked DNA. To test whether bypass was affected by position of the roadblock we created a dCas9 block at target site A which is located close to GATC site 1. No change in roadblock bypass efficiency was observed (Supplemental Figure S2). Taken together, despite placement of a large, specific and stable dCas9 RNP complex in between mismatch and GATC site, considerable MMR initiation takes place beyond the roadblock, similar to the reported GATC site incision beyond the EcoRI_E111Q_ roadblock ^31^.

### A functional single molecule assay for MMR strand incision

To further establish the ability of MMR components to bypass a roadblock during strand incision, we implemented a series of single-molecule assays. First, we designed a ~3-kbp DNA construct based on the pREP4 plasmid with a single-base (dT) insertion flanked on either side (at 270 and 570 bp from the insertion) by an unmethylated GATC incision site (see Material and Methods for details). Using a magnetic trap to supercoil this DNA allowed generation of interwound “plectonemic” structures which reduce the DNA extension (Figure 2A). Approximately 90-95% of DNA molecules in the magnetic trap can be supercoiled in this manner. In the presence of 2.5 nM MutS, 10 nM MutL, 10 nM MutH, and ATP, incision of a GATC site on this DNA is observed as an abrupt return of DNA extension to the initial, unsupercoiled state due to rapid relaxation of plectonemic supercoils (Figure 2B). The presence of T4 DNA ligase in the reaction allows the nicked DNA to be ligated and readily be supercoiled anew. Typically, on the order of 80% of DNA molecules which can be supercoiled in the magnetic trap undergo repeated cycles of incision. Control experiments underscore the specificity of this reaction, as replacing the heteroduplex DNA with homoduplex DNA, methylating the GATC sites, or withholding any one of the protein components or ATP, completely abolishes the incision reaction (Supplemental Figure S3). Determination of T_wait_, the time between formation of DNA supercoils and incision, shows that this variable is exponentially distributed for a mean lifetime of the supercoiled state <T_wait_> = 96 ± 11 s SEM (n = 150, Figure 2C). This assay establishes quantitative single-molecule detection of DNA incision by the wild-type *E. coli* MMR system.

**Figure 2:**
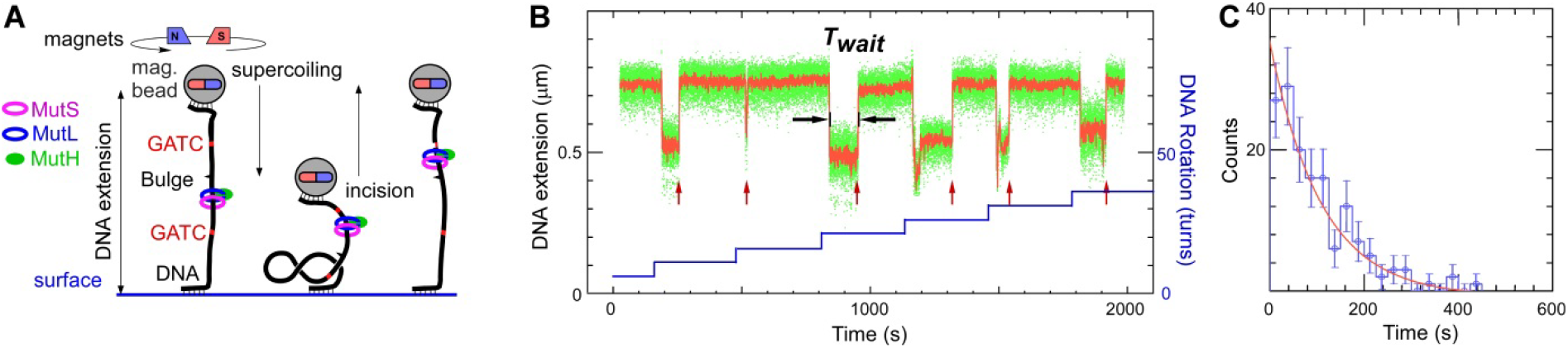
Single molecule DNA nanomanipulation assay for MMR strand incision. A) The heteroduplex pREP4 DNA substrate is tethered and supercoiled in the magnetic trap in the presence of MMR components. Incision of DNA is observed as an abrupt increase in DNA extension. B) Time-trace showing repeated cycles of supercoiling and subsequent incision of the substrate by 2.5 nM MutS, 10 nM MutL, 10 nM MutH. In between cycles incised DNA is religated by T4 DNA ligase which is present in the reaction. Green points show bead position sampled at 30 Hz, red points show raw data with ~2 s averaging. Blue line represents the stepwise increase in supercoiling imposed on the DNA via the magnetic trap. Red arrows indicate incision events. T_wait_ represents the time elapsed before a supercoiled DNA is incised. C) Distribution of T_wait_ is well-described by a single-exponential fit (red line) with mean of 96 ± 11 s (SEM, n=150).

### Bypass of a stalled RNA polymerase roadblock

Using this single-molecule assay we then tested the ability of the MMR components to bypass a roadblock that is observable, reversible and larger than the dCas9-RNA complex. For this we designed a DNA construct in which *E. coli* RNA polymerase (RNAP, 450 kDa) could be initiated and stalled on a so-called “C-less cassette” positioned between a mismatch and an unmethylated GATC site (Rb-pREP4 construct, see Materials and Methods). The stalled transcription elongation complex (sTEC) is stable on the nanomanipulated DNA and its formation and presence can be continuously monitored in the magnetic trap ^39^. In addition, the sTEC can be quantitatively chased from the DNA by addition of GTP and confirmed to be absent using the magnetic trap ^39^.

We first characterized DNA strand incision on the Rb-pREP4 construct in the absence of RNAP using 2.5 nM MutS, 10 nM MutL and 10 nM MutH, see Figure 3A; Table 1. Once again, ~80% of DNA molecules which can be supercoiled undergo multiple rounds of incision by the MMR components, and the mean lifetime of the supercoiled state prior to incision is found to be 186 ± 11 s SEM (n=590 incision events collected from 93 DNA molecules; Table 1).

**Table 1.** Statistics of the number of molecular events monitored during single-molecule roadblock experiments.

|  | MutS, MutL, MutH |  | MutS, MutLΔ4, MutH |  |
| --- | --- | --- | --- | --- |
|  | No RNAP | Stalled RNAP | No RNAP | Stalled RNAP |
| # of DNA molecules monitored | 115 | 208 | 48 | 108 |
| # of DNA molecules with stalled RNAP |  | 117 |  | 90 |
| # of DNA molecules incised | 93 | 20 | 38 | 0 |
| Mean Twait | 186 ± 11s (SEM, n=590) | 530 ± 73s (SEM, n=187) | 666 ± 233s (SEM, n=206) |  |

**Figure 3:**
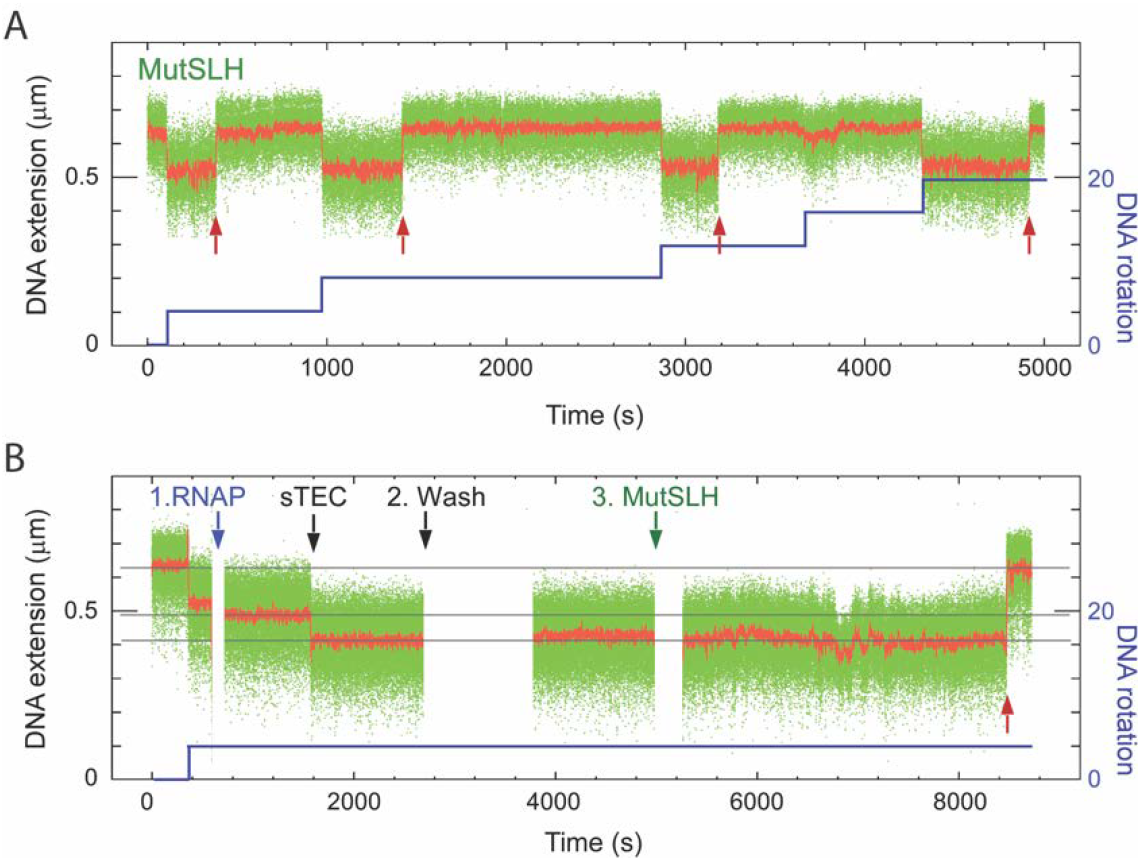
Effect of RNAP roadblock on single molecule strand incision. Time-traces of incision assays carried out on the Rb-pREP4 construct using 2.5 nM MutS, 10 nM MutL, and 10 nM MutH (**A**) in the absence or (**B**) in the presence of stalled RNA polymerase (RNAP). Red arrows show incision events, blue line the supercoiling pattern imposed in the magnetic trap. RNAP addition (**1**) is rapidly followed by formation of a stalled elongation complex (sTEC). (**2**) Free components are washed out and (**3**) MMR components introduced. Incision is observed when wild-type MMR components are used despite the presence of the roadblock (B, red arrow).

When we stalled RNAP thirty bases from the transcription start site using the C-less cassette and then added the same MMR components as above, DNA incision was severely – but not completely – inhibited (Figure 3B; Table 1). Indeed the fraction of DNA molecules which could be incised is now only 17% (20 out of 117 DNA molecules), and the mean lifetime of the supercoiled state observed on those molecules which did display incision increased to 530 ± 73 s SEM (n=187 incision events collected from 20 DNA molecules, Table 1). This confirms that MMR components possess the ability to functionally bypass large, fixed roadblocks.

### Shortening of MutL linker length abolishes roadblock bypass

To further investigate roadblock bypass, a MutL variant lacking amino acid 366 to 405 (MutLΔ4, ^40^) was used to replace wild type MutL in the incision assays. In this deletion mutant, the disordered linkers in between the N- and C-terminal globular domains in both subunits of the MutL dimer have been shortened from 100 to 60 residues. MutLΔ4 has a smaller hydrodynamic volume when compared to wild-type MutL both in the nucleotide-free, open state as well as in the nucleotide-bound, closed conformation (Supplemental Figure S4;^40^). In the absence of a dCas9 roadblock, MutLΔ4 behaved similar to wildtype MutL in the ensemble incision reaction (Figure 4A). However, in the presence of a dCas9 roadblock on target site C in between the GATC sites on substrate GT#2, exclusively GATC site 2 close to the mismatch was incised. No reaction intermediates or products containing an incision at GATC site 1 located beyond the dCas9 roadblock were observed. Correspondingly, nicking beyond the roadblock was also abolished on the GT#2b substrate (Figure 4B).

**Figure 4:**
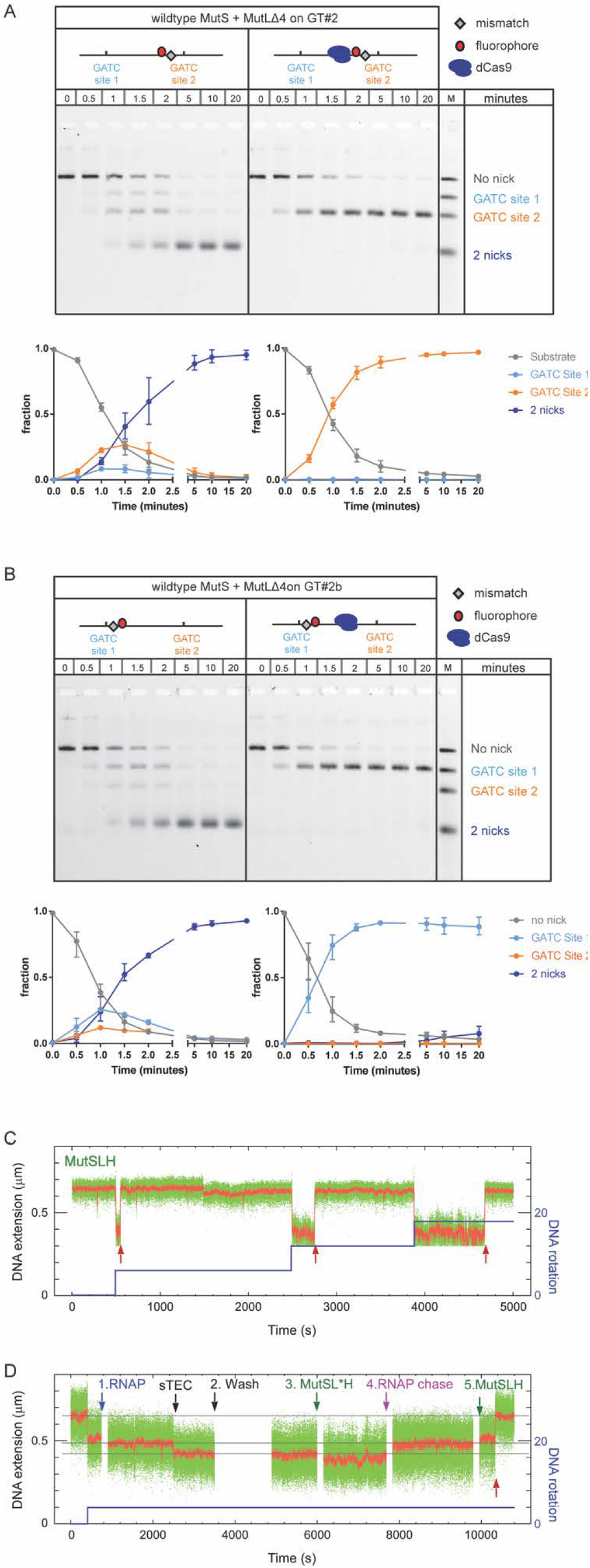
Shortening of MutL linker regions prevents roadblock bypass. Time courses for GATC site incision by 50 nM MutS, 50 nM MutLΔ4 and 25 nM MutH on 0.5 nM GT#2 substrate (panel A) and 0.5 nM GT#2b (panel B) in the absence and presence of a dCas9 roadblock as indicated. Reaction products are separated using gel electrophoresis under denaturing conditions and visualized using the Alexa^647^ fluorophore. Graphs show quantification of product fractions containing no nick (gray), a nick at GATC site 1 (light blue), a nick at GATC site 2 (orange) and 2 nicks (dark blue). Data points with error bars represent the mean values and range of three independent experiments. Time-traces of incision assays carried out on the Rb-pREP4 construct using 2.5 nM MutS, 10 nM MutL∆4, and 10 nM MutH (**C**) in the absence or (**D**) in the presence of stalled RNAP. Red arrows show incision events, blue line the supercoiling pattern imposed in the magnetic trap. RNAP addition (**1**) is rapidly followed by formation of a stalled elongation complex (sTEC). (**2**) Free components are washed out and (**3**) MMR components introduced. Incision is observed when wild-type MMR components are used despite the presence of the roadblock (B, red arrow). The roadblock prevents incision when MutL∆4 is used instead of MutL, but when (D**4**) the roadblock is lifted and (D**5**) MutS, MutL∆4 and MutH are introduced, incision resumes (red arrow).

The single-molecule incision and roadblock assays confirm these results. On the Rb-pREP4 construct the combination of 2.5 nM MutS, 10 nM MutLΔ4, and 10 nM MutH generated a robust reaction, with 80% of supercoilable DNA molecules undergoing incision with a mean waiting time of 666 ± 233 s SEM (n=206 events collected from 38 DNA molecules, see Figure 4C and Table 1). When RNAP was stalled on the CTP-less cassette prior to addition of the same MMR components, strictly no incision was observed on any of the 90 DNA molecules bearing stalled RNAP (n=0 events collected from 90 DNA molecules monitored, see Table 1). Chasing off the roadblock and reintroduction of MutS, MutLΔ4 and MutH restored incision (Figure 4D).

Thus, our combined functional ensemble and single molecule analyses reveal that shortening the length of the disordered MutL linkers completely abolishes roadblock bypass during MMR strand incision.

## Discussion

In this study we address the mechanism of communication between mismatch and strand discrimination signal during MMR daughter strand incision. Placing a roadblock in between mismatch and GATC site allowed us to distinguish between mechanisms that signal along the DNA backbone and through three-dimensional collision. While the former should be inhibited by a block on the DNA, the latter should not be affected to the same extent. Our observation that roadblock bypass can be fully attributed to the long, disordered linker regions connecting the globular N-and C-terminal domains in the MutL protein establish that communication during MMR initiation occurs along the DNA backbone. Below we discuss the implications of these findings for DNA MMR as well as for other complex biochemical reactions that take place in crowded environments.

### Quantitative assays that monitor incision at individual GATC sites and on single DNA molecules

We adapted our ensemble incision assay ^35^ such that we could quantitatively monitor incision of individual hemimethylated GATC sites that flank the mismatch on substrates GT#2 and GT#2b ^35^. The estimated maximal accumulation of reaction intermediates that are nicked either at GATC site 1 or GATC site 2 adds up to 30% of total substrate. This is similar to the observed amount of single-nicked intermediate obtained on circular DNA at otherwise identical conditions (see Figure 3C left panel in ^35^); indicating that also on linear DNA dual GATC site incision is processive ^35^. Interestingly, while no GATC site preference is observed on circular substrates, linear substrates reveal clear preference for the GATC site closest to the mismatch. Site preference thus depends on the configuration of the DNA substrate and is not an effect of the sequence context of a particular GATC site. The capability to observe GATC site preference emphasizes the resolving power of this new assay.

We furthermore established a functional single molecule assay for DNA MMR. Using DNA nanomanipulation we could monitor incision of a GATC site by the MutSLH system in real time on individual mismatch-containing DNA molecules. Control experiments demonstrate that incision is absolutely specific to mismatch-bearing DNA and required all three protein components as well as ATP. In the presence of T4 DNA ligase DNA molecules could be ligated and incised numerous times, allowing us to obtain a direct quantitation of the rate of incision.

### Nuclease-deficient Cas9 and RNA polymerase as versatile in vitro roadblocks

In this work we made use of two *in vitro* protein blocks. *Streptococcus pyogenes* Cas9 is a multidomain endonuclease that can be guided by RNA to bind to specific DNA target sites ^36,41^. As expected ^36-38^, we found that dCas9 forms a specific and stable complex on different target sites and remains bound for many hours. We have previously used the D10A variant of Cas9 as *in vitro* endonuclease to create substrates to analyze strand excision in a reconstituted MMR system ^35^. An advantage of using Cas9 and its variants in this kind of approaches is that PAM sites are ubiquitously present in DNA and thus target site selection merely requires addition of a suitable crRNA, rather than requiring cloning procedures to introduce or change the location of binding sites. Specifically, in our ensemble assay this allowed us to easily change the location of the dCas9 block to determine the effect of the distance between the block and individual GATC sites.

In our single molecule incision assay we generated a controllable, observable and reversible roadblock by stalling *E. coli* RNA polymerase in between the mismatch and the GATC site. Like the Cas9-RNP complex, RNAP relies on protein-DNA as well as DNA-RNA base-pair formation to form a stable and specific complex on a target site. RNAP (~450 kDa) forms a considerably larger roadblock than EcoRI_E111Q_ (62 kDa) or the dCas9-RNP complex (~ 190 kDa). Because the continuous presence of the stalled RNAP throughout the course of the single-molecule incision assays could be verified through its well-established signature in nanomanipulation assays ^39^, we were able to ascertain that the roadblock protein was not microscopically vacating its recognition site. Furthermore, because RNAP is stalled at its promotor by withholding CTP, the roadblock is reversible. Addition of the missing nucleotide chases off RNAP, which allows restoration of incision activity thus confirming that the observed effect on nicking efficiency indeed was caused by the presence of the block.

### Roadblock bypass during MMR incision

One of the few studies combining behavior of MMR proteins on DNA with functional readout ^31^ reports that placement of an EcoRI_E111Q_ roadblock in between a mismatch and a single GATC site on linear DNA inhibits GATC site incision. However, inhibition was incomplete, with approximately 30% activity remaining compared to activity in the absence of a roadblock. The authors attributed residual incision activity beyond the roadblock to the block temporarily vacating its specific binding site, MMR proteins being able to bypass the block through loop formation ^40,42^, or both of these mechanisms. Likewise, we observed roadblock bypass in our analyses; despite being larger than EcoRI_E111Q_, both the dCas9 and RNAP roadblocks reduce the rate of incision of the GATC site beyond the block, but do not completely inhibit it. Because the residence time of dCas9 on a target site is considerably longer than for EcoRI_E111Q_ (>6 hours compared with ~40 minutes) and because stalled RNAP unequivocally remains bound to the C-less cassette while the GATC site beyond this complex gets incised, residual GATC site incision in our assays is not due to roadblocks microscopically vacating their binding site.

Concerning loop formation, the observed preference for the GATC site close to the mismatch in our ensemble incision assay does not correlate with the DNA cyclization probability (the ‘J-factor” distribution) that governs thermal loop formation ^43^; this distribution predicts that loop formation would be most probable between sites approximately 1 kb apart (which in our system would be the GATC site far away from the mismatch) and unlikely at distances below 100 bp (GATC site close to the mismatch). Likewise, Modrich and colleagues observed complete inhibition of incision upon introduction of a double strand break between the GATC site and the mismatch, while on this substrate thermal loop formation should still have allowed incision by bringing mismatch and GATC site in close proximity ^31^. We thus considered an alternative explanation for observed GATC site incision beyond a roadblock: the reported ability of the MutL protein to bypass MutS sliding clamps ^32^ and its eukaryotic homolog MutLα to bypass MutSα, other MutLα molecules and nucleosomes ^33^ while diffusing along DNA.

### Roadblock bypass is enabled by long disordered linkers of MutL

The MutL protein is a homodimer in prokaryotes, and a heterodimer in eukaryotes ^1^. In both cases, each subunit contains N- and C-terminal globular domains connected by a long, disordered linker. In *E. coli* MutL these linkers are 100 amino acids in length whereas yeast homologs Mlh1 and Pms1 linker arms contain approximately 150 and 250 amino acids, respectively ^40,44^. MutL undergoes conformational changes in the presence of nucleotide cofactors, particularly ATP ^35,40,45-47^. Upon ATP binding, the N-terminal ATPase domains of *E. coli* MutL dimerize such that the protein forms a ring-like structure which encloses DNA ^32,40^. In eukaryotic MutLα the conformational changes involve condensation of the flexible linkers, thereby bringing together the globular domains ^47^. Cleavage of the yeast MutLα linkers impairs *in vivo* MMR, indicating that the linkers must be intact ^44^. Deletions in the Mlh1 protein are tolerated less well than in Pms1 ^44^. The *E. coli* MutLΔ4 variant, in which the linker length has been reduced by 40%, is fully functional in our ensemble incision assay, both on GATC sites close to and far away from the mismatch. In the presence of a roadblock, the complete absence of GATC site incision beyond the blocks in both our assays with MutLΔ4 convincingly shows that bypass can be fully attributed to the long linker regions in MutL. Because MutLΔ4 is fully functional on DNA without a block, loss of incision at the site beyond the block is due to the block itself and not due to an effect of the deletion on the capability of MutLΔ4 to communicate to GATC sites far away.

### Implications for the mode of communication during MMR initiation

Over the years, multiple models have been proposed for the molecular mechanism that enables communication between mismatch and GATC site during initiation of MMR ^20^. These involve ATP-driven active loop extrusion by MutS ^48,49^, static transactivation by formation of protein-mediated loops from mismatch to GATC site 42,50, polymerization ^51^ in which filaments of MutL along the DNA backbone connect MutS at the mismatch to the strand discrimination site ^52,53^ and the molecular switch model in which the proteins diffuse in an ATP-hydrolysis independent manner along the DNA ^8,14^. Growing support for models that describe communication along the helix contour has recently been provided by single-molecule analyses in which *E. coli*, *Thermus aquaticus* and yeast MutS homologs as well as *E. coli* and yeast MutL homologs have been shown to diffuse along DNA after mismatch recognition in the presence of ATP ^32-34^. However, these observations were not directly coupled to functional analysis, and therefore it is not clear whether the observed diffusion relates to actual strand incision.

Our work establishes that roadblock bypass during strand incision is due to the globular domains of MutL being connected with long disordered linkers. This finding further argues against thermal loop formation being responsible for bringing together mismatch and GATC site. The MutLΔ4 variant efficiently incises the GATC site that is far away from the mismatch in the absence of a roadblock, and is thus not affected in its capability to communicate. If loop formation would drive incision (Figure 5A), MutLΔ4 should still be able to incise the GATC site even in the presence of a roadblock. However, placement of a block fully inhibits incision upon shortening the unstructured MutL linkers. Thus, the loss of incision is solely due to placement of the block. Therefore, communication between mismatch and GATC site during MMR initiation occurs exclusively via the DNA helix contour. Whether the mechanism of this communication along the DNA backbone involves diffusion ^8,14,32^, active loop extrusion ^48,54^ or polymerization ^51,52^ remains to be established.

**Figure 5:**
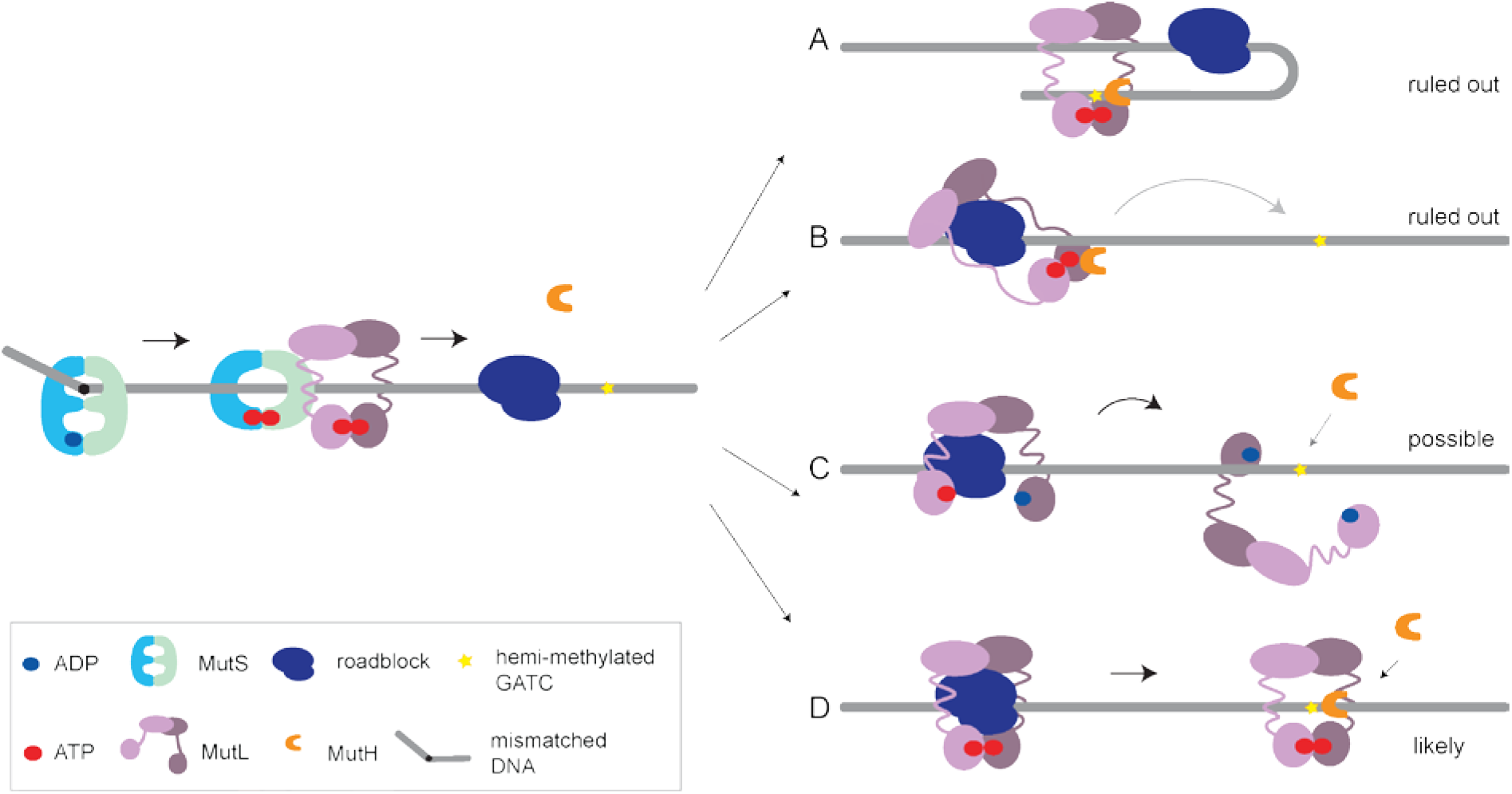
The long unstructured linkers of MutL enable roadblock bypass during MMR initiation. Mismatch recognition by MutS results in binding of ATP and conversion into the activated sliding clamp conformation. This results in binding of MutL which is then activated by ATP and can recruit MutH to incise a hemimethylated GATC site. Upon encountering a roadblock on the DNA, bypass can occur as follows: A) Activation *in trans* through loop formation or strand transfer. This is ruled out because incision mediated by MutLΔ4, which occurs in the absence of a roadblock, would not be completely inhibited via this mechanism by presence of a block further along the DNA. B) The disordered linkers allow one region of MutL to reach over the roadblock towards the GATC site while remaining in its activated state. This mechanism is ruled out as incision of GATC sites that are beyond the reach of the fully extended linker regions has been observed. C) The disordered linkers allow MutL to hop over a roadblock *via* temporarily adopting an open state. This mechanism will only allow GATC site incision by MutH if after the hop MutL is still in its activated state. Because activation of MutL involves MutS-mediated loading and subsequent closure, and because MutS cannot bypass roadblocks, it is unclear how MutL that hops *via* ATP hydrolysis and opening at its N-terminal domains would be able to activate MutH beyond the roadblock. Hopping facilitated through temporarily opening at the C-terminal domains would possibly maintain the active state. D) The disordered linkers create an internal MutL cavity with a size that is large enough for MutL to move over roadblocks on DNA while remaining in its activated state, thereby being able to support MutH-mediated GATC site incision beyond the block. Shortening the linker regions in MutLΔ4 results in a smaller cavity that can no longer thread over roadblocks.

The 100 amino acids-long linkers in MutL are disordered and because they are proline-rich, are expected to be relatively extended due to the restricted backbone torsion angles ^40,55^. It is therefore possible that while one globular region of MutL remains bound on the mismatch-proximal site of the roadblock, the long linkers allow reaching out over the block allowing the other globular region of MutL to activate MutH on a GATC site at the mismatch-distal site of the block (Figure 5B). We find this mechanism unlikely because the distance between the roadblock and the hemimethylated GATC does not influence bypass (Supplemental Figure S1) and the length of the linkers (350 Å each in fully extended state) is not long enough to reach over a block to the far GATC site that is located 800 bp (~2800 Å) from the block.

An attractive mechanism for roadblock bypass is the behavior observed for eukaryotic MutLα ^33^ during which the protein may transiently adopt an open ring configuration to hop over a roadblock (Figure 5C). If hopping indeed takes place, then it is not clear whether the transient opening is correlated with hydrolysis of bound ATP in the N-terminal domains and how, in the absence of MutSα beyond the roadblock, MutLα then would retain or would be able to reform its active conformation. Future experiments using cross-linked MutL or variants defective in opening might shed further light on this mechanism.

A second attractive mechanism is diffusion of MutL over the block (Figure 5D). The estimated diameter of the central cavity encircled by the disordered linkers is ~100 Å based on observed Stokes radii ^40^. This diameter would be large enough to allow a DNA bound EcoRI_E111Q_ (size ~75 Å) to pass through the ring, as well as the Cas9 RNP complex (size ~100 Å). As the linkers also have been shown to contain flexible regions ^47^ only a minor expansion would be required to allow RNAP (130-150 Å) to pass through (fully extended linkers are estimated to be able to form a ring with ~200 Å diameter). Thus, simple diffusion of MutL sliding clamps over a roadblock would allow the observed bypass (Figure 5D).

The observed roadblock bypass has interesting implications for the composition of the functional incision complex during initiation of MMR. While it is evident that MutL can bypass blocks, available data suggest that MutS is unable to do this. If MutS sliding clamps would be able to bypass the block in our functional assay, it would be expected to bind MutL and activate incision beyond the block, which we do not observe for the MutLΔ4 variant. Furthermore, yeast homolog MutSα is unable to bypass barriers on DNA ^33,56^. Indeed, the size of the central cavity in the MutS sliding clamp (~25 Å; ^10^) is just large enough to accommodate a DNA duplex but too small to allow any of the roadblocks to pass through. MutL separates from MutS and diffuses along DNA on its own or in complex with MutH ^32^. Taken together these observations suggest that MutH-mediated GATC site incision beyond the roadblock is initiated by MutL that has released MutS and has physically bypassed the roadblock. MutS therefore is not necessarily part of the actual strand incision complex that nicks the DNA ^32^.

*In vivo*, processes on DNA take place in a crowded environment. Even when the DNA replicative machinery and its associated proteins temporarily evict bound proteins from DNA during replication ^57^, protein complexes will reassemble behind the replication fork and may interfere with the search for the strand discrimination signal following mismatch detection. In *E. coli,* roadblock bypass would allow MMR to continue despite proteins like H-NS and HU being bound to the DNA. Based on analysis of one specific locus, the *rpob* gene, the MutLΔ4 mutant does not display a mutator phenotype ^40^, and one could, therefore, conclude that roadblock bypass does not have to occur very often. However, the *rpob* gene is heavily transcribed and rich in GATC sites, and the chance of hitting a roadblock at this locus may therefore be lower than at a silent locus that is devoid of GATC sites (meaning longer stretches of DNA have to be scanned after mismatch recognition). In eukaryotes, MutSα delays CAF1-mediated nucleosome repositioning ^58^, nevertheless any nucleosome that is (still) present can be bypassed by MutLα. If indeed, as in the *E. coli* system, the MutLα is still functional after bypassing the block, this could also be the mechanistic explanation for the observed initiation of excision on DNA containing streptavidin blocks in human cell extracts ^59^.

Proteins with intrinsically disordered regions are remarkably abundant in all kingdoms of life ^60^. While sequence conservation is not always very high in these regions, they nevertheless serve specific and often essential roles in biological processes. Unstructured regions have for example been identified in proteins implicated in cell signaling, transcription, chromatin remodeling and DNA repair ^45, 61-63^. All these processes have to take place in the crowded cellular environment. One general role of unstructured regions may be that they provide proteins with the capability and/or flexibility to work around challenges provided by these crowded environments. As shown here, this may be particularly relevant for processes that occur on DNA or other cellular structures such as microtubules in which communication takes place in one-dimensional space.

In conclusion, we have shown here that MMR proteins communicate along the DNA helix contour during DNA strand incision, and that previously observed roadblock bypass can be fully attributed to the long, disordered linkers in MutL. The two new assays that we have developed allow further unravelling of the molecular mechanism that is used by MMR proteins to communicate along the DNA helix, and opens the way for quantitative, bottom-up reconstruction of the complete MMR pathway at single molecule resolution.

## Online Methods

### Protein purification

*E. coli* MutS, MutL, MutH, RNA polymerase and σ^70^ transcription factor were purified and stored as described previously ^10, 64-67^. The expression construct for MutLΔ4 lacking amino acids 366-405 ^40^ was created using Quikchange mutagenesis on wild-type MutL expression plasmid pTX417 (forward primer 5’-GGGGCGCAATCACTTTGCAGAACGCCAGCTTTTGCAAACGC; reverse primer 5’-GCGTTTGCAAAAGCTGGCGTTCTGCAAAGTGATTGCGCCCC). The resulting construct was sequence verified and transformed into BL21(DE3),mutL::Tn10 (derived from strain BL21(DE3)pLysS,mutL::Tn10 (gift from N. Goosen) through five-fold serial dilution in LB containing 100 μg/ml acridine orange). MutLΔ4 was purified using the same procedure as for wild type MutL. Protein concentrations were determined spectrophometrically (ϵ^280 nm^ =73605 M^-1^ cm^-1^ for MutS, ϵ^280 nm^ =54270 M^-1^ cm^-1^ for MutL, ϵ^280 nm^ =44920 M^-1^ cm^-1^ for MutLΔ4, ϵ^280 nm^ =38023 M^-1^ cm^-1^ for MutH). Cas9 containing amino acid substitutions D10A and H840A (dCas9) was purified from Rosetta2 (DE3) pLysS cells carrying expression construct pMJ841 (gift from Jennifer Doudna; Addgene plasmid # 39318) according to the following procedure: dCas9 was expressed over night at 16°C in LB containing 25 μg/ml kanamycin and 33 μg/ml chloramphenicol using 1 mM isopropyl 1-thio-β-D-galactopyranoside (IPTG). Cells were harvested, resuspended in buffer A (20 mM Tris pH 8.0, 500 mM NaCl, 20 mM imidazole, 2 mM β-mercaptoethanol) containing 1 mM phenylmethylsulfonyl fluoride (PMSF) and cOmplete EDTA-free protease inhibitors (Roche Diagnostics GmbH, Mannhein, Germany) and lysed by sonication. Cleared supernatant was mixed with Nickel Sepharose 6 Fast Flow (GE Healthcare Biosciences, Uppsala, Sweden). Beads were washed with buffer A containing 0.1 mM PMSF and subsequently with buffers containing 1 M NaCl and 250 mM NaCl. dCas9 was eluted using 20 mM Tris pH 8.0, 250 mM NaCl, 10% glycerol, 250 mM imidazole, 2 mM β-mercaptoethanol. Eluate was adjusted to 1 mM DTT and 0.5 mM EDTA, Tobacca Etch Virus (TEV) protease was added and the mixture was dialyzed over night at 4°C to remove the His-MBP tag against buffer B (25 mM Hepes pH 7.5, 2 mM β-mercaptoethanol, 10% glycerol) containing 150 mM KCl. Dialysate was brought to 20 mM imidazole, incubated for 30 minutes with Nickel Sepharose beads to remove TEV and uncleaved dCas9. dCas9 was further purified using chromatography on HiTrap SP sepharose, heparin and Superdex 200 16/60 columns (GEHealthcare) in buffer B containing appropriate KCl concentrations for each step. dCas9 aliquots in 25 mM Hepes pH 7.5, 150 mM KCl, 10 mM β-mercaptoethanol and 10% glycerol were flash-frozen and stored at −80°C. dCas9 protein concentration was determined spectrophotometrically (ϵ^280 nm^ = 120450 cm^-1^ M^-1^).

### DNA constructs for ensemble incision assay

Hemimethylated DNA substrate containing a single G/T mismatch or the corresponding A/T Watson Crick base pair were generated by primer extension using a primer containing an Alexa Fluor ^®^ 647 (Alexa^647^) fluorophore (IBA GmbH, Göttingen, Germany) on single stranded DNA derived from phagemid GATC2 ^35^. Closed circular DNA was purified from gel using the Wizard® SV Gel and PCR Clean-Up System (Promega, Madison, USA). Primer GT14 [5’P-CCAGACGTCTGTC-g-ACGTTGGGAAGCTT*GAGTATTCTATAGTGTCACCT 3’], where ‘g’ indicates the guanidine forming the mismatch and ‘T*’ is the Alexa^647^-labeled nucleotide,was used to construct GT#2^647^ substrate (two GATC sites 31 bp 5’ and 1042 bp 3’ of the mismatch). Primer AT14 [5’P-CCAGACGTCTGTCTACGTTGGGAAGCTT^*^GAGTATTCTATAGTGTCACCT 3’] was used to construct the homoduplex substrate. Primer GT28 [5’P-GGTAGCTCTTCAT*CCGGCAAACAAACC-g-CCGCTGGTAGCG] was used to construct the GT#2b^647^ substrate (two GATC sites 61 bp 3’ and 1012 5’ of the mismatch). DNA substrates were linearized by ScaI-HF (New England Biolabs) for 30 min at 37°C followed by heat inactivation for 20 min at 70°C. Marker was created as follows: GT#2^647^ substrate was linearized with ScaI-HF to create the full-length linear fragment. The fragment indicative for nicking at GATC site 1 was created by digesting GT#1b ^35^ for 30 min at 37°C with ScaI-HF followed by 30 min at 60°C with BstYI. The fragment indicative for nicking at GATC site 2 was created by digesting GT#2^647^ using ScaI-HF and BamHI for 30 min at at 37°C. The fragment indicating 2 nicking events was prepared by digesting GT#2^647^ with BstYI for 30 min at 60°C. Fragments were mixed at equimolar ratio.

### DNA constructs for single-molecule nanomanipulation

#### pREP4

The DNA construct used to establish the single-molecule incision assay is based on the ~3 kbp XbaI-SacI fragment of the pREP4 plasmid (Invitrogen). This fragment contains a unique SpeI site flanked on one side by a 5’-GATC site located 570 bp away and on the other side by a 5’-GATC site located 270 bp away. Double-strand DNA oligos either heteroduplex or homoduplex and containing phosphorylated XbaI overhangs were ligated overnight and at room temperature into the SpeI site of the vector. This was carried out in a final volume of 160 μl of 1x CutSmart buffer (New England Biolabs) in a 3-fold molar excess of insert over vector (3 μM and 1 μM, respectively) using 6000 units of T4 DNA ligase (New England Biolabs) and in the additional presence of 50-100 units of SpeI-HF restriction enzyme (New England Biolabs). Heteroduplex oligo was obtained by annealing primers MutS0-Bot (5’P-CTAGACTGAAGACCTTCTGCCGGCCGTCTGGACCTGTCT) and MutS1-Top (5’P-CTAGAGACAGGTCCAGACGGCACGGCAGAAGGTCTTCAGT) which differ in length by one base. Homoduplex oligo was obtained by annealing primers MutS0-Bot and MutS0-Top (5’P-CTAGAGACAGGTCCAGACGGCCGGCAGAAGGTCTTCAGT). Subsequent cleavage of the ligation product with XbaI, SacI and SpeI site generated a 3 kb fragment containing one and only one insert of the heteroduplex or homoduplex oligo as well as an XbaI overhang and a SacI overhang. This 3-kb fragment was gel purified and ligated to 1-kb biotin-labeled DNA bearing an XbaI overhang and 1-kb digoxigenin-labeled DNA bearing a SacI overhang as previously described ^68^.

#### Rb-pREP4

A DNA construct containing a transcription cassette was developed to test MMR incision past a stalled RNAP roadblock. The GATC sites contained within the XbaI-SacI fragment of pREP4 were removed using site-directed mutagenesis, and the native SpeI site was repositioned 300 bp away from the SacI site. Next a transcription cassette was inserted 350 bp downstream of the SpeI site, and a single GATC site 350 bp from the promoter of the transcription cassette. The transcription cassette contains the early N25 bacterial promoter from phage T5 (for which the first thirty bases of transcript require only ATP, UTP and GTP to be transcribed) and a 71-bp transcribed region followed by a bacterial tR2 transcription terminator ^39^. Thus, in this construct a stalled RNAP provided only ATP, UTP and CTP forms a roadblock roughly halfway between the mismatch-insertion SpeI site and the GATC site. The stalled RNAP can be chased from the construct by adding GTP to the reaction. Heteroduplex oligo was incorporated into this vector as described above, and the construct obtained was derivatized with biotin-labeled DNA and digoxigenin-labeled DNA as described above.

### Creation of dCas9 roadblocks on mismatch-containing hemimethylated DNA substrates

Alt-R TracrRNA and crRNA (rCrUrU rCrUrA rGrUrG rUrArG rCrCrG rUrArG rUrUrG rUrUrU rUrArG rArGrC rUrArU rGrCrU for block A, rUrGrU rGrArG rUrUrA rGrCrU rCrArC rUrUrG rUrUrU rUrArG rArGrC rUrArU rGrCrU for block C) were purchased from IDT (Leuven, Belgium). Duplex RNA was formed by incubating a mixture containing 3 μM of tracrRNA and 3 μM of the appropriate crRNA oligonucleotide in IDT duplex buffer (IDT, Leuven, Belgium) for 5 min at 95°C and slow cooling at room temperature. The dCas9-RNA complex was formed by incubating 80nM dCas9 with 600nM duplex RNA in 25 mM Hepes KOH [pH 7.5], 150 mM KCl, 5 mM MgCl_2_, 1 mM DTT, 10% glycerol, 100 ng/μl BSA, 0.005% Tween 20 for 5 min at 37°C. The ribonucleoprotein complex was incubated with GT#2 or GT#2b linear DNA substrate for 5 min at 37°C in a final concentration of 20 nM dCas9 (and thus 150 nM RNA) for 0.5 nM DNA. Block C is positioned in the center of the substrate at 148 bp and 813 bp away from the mismatch in GT#2 and GT#2b, respectively. Block A is positioned 879 bp and 82 bp away from the mismatch in GT#2 and GT#2b, respectively. These distances are large enough for MutS binding to the mismatch to not be obstructed by the blocks.

### MutH activation assay

Strand incision assays were performed with 50 nM MutS, 50 nM MutL or MutLΔ4, 25nM MutH and 0.5 nM Alexa^647^ labelled linear DNA substrate at 37°C in standard assay buffer (25 mM Hepes KOH [pH 7.5], 150 mM KCl, 5 mM MgCl_2_, 1 mM DTT, 10% glycerol, 100 ng/μl BSA, 0.005% Tween 20) containing 1mM ATP. Reactions were started by adding a mixture of DNA and ATP to a mixture of MutS, MutL and MutH. Reactions (10 μl) were stopped at the indicated time points by adding 40 μl stop solution containing 8M Urea and 0.3% SDS. Samples were denatured by incubation for 10 min at 85°C and run on a 1.5% agarose gel in 1^*^TBE, both gel and buffer containing 40 μM chloroquine and 2M urea. Incision product formation separated by gel electrophoresis was monitored using a Typhoon FLA imager (GE Healthcare). The Alexa^647^ fluorophore was excited at 633nm and emission was passed through the Cy5 670BP30 filter. Band intensities were quantified using FIJI software ^69^. Fractions of substrate without nick, nicked intermediates and products containing two nicks were calculated relative to the total signal obtained from the fluorophore.

### Single-molecule incision and roadblock assays and data analysis

Magnetic trapping was used to nanomanipulate individual DNA molecules. Briefly, in the magnetic trap the ~3-kbp DNA substrate (~1 μm contour length) is tethered to a glass surface at one end and to a 1-μm superparamagnetic bead at the other. A pair of magnets located above the sample is used to gently extend the DNA away from the surface; the extending force can be increased or decreased by, respectively, moving the magnets closer to or farther away from the glass surface. The DNA can also be supercoiled by rotating the magnets. At the low extending force used here (F ~ 0.25 pN; 1 pN = 10^-12^ Newtons) supercoiled DNA forms plectonemic loops, and extension of a plectonemic loop by a single topological unit of writhe causes the DNA extension to decrease by a fixed length (here typically ~60 nm; ^70^). On supercoiled DNA, single-strand incision is detected as an abrupt increase in DNA extension as plectonemes are essentially instantly dissipated.

DNA tethers for nanomanipulation were assembled as follows. Biotin- and digoxigenin-labeled DNA constructs for single-molecule experiments were diluted to 10-50 pM in 10 mM Tris buffer (pH 8). Five microliters of superparamagnetic beads (Dynal MyOne C1, Life Technologies) were washed in reaction buffer RB (20 mM K∙Hepes pH 7.5, 150 mM KCl, 5 mM MgCl_2_, 0.05% Tween-20, 1 mg/ml BSA, 1 mM DTT) and resuspended in ten microliters of RB. The beads were then combined with one microliter of DNA and left to incubate for a few seconds before the reaction was gently diluted by addition of a further 20 μl of RB.

The bead-DNA mixture was introduced into a 10-μl flowcell consisting of two #1 glass coverslips coated with polystyrene, separated by two layers of parafilm with a channel cutout, and functionalized with antidigoxigenin as previously described ^68^.

Solutions can be added or removed from this flowcell at will via appropriate inlet and outlet ports. After beads have been left to sediment for 5 minutes the chamber is washed with 2-5x 500 μl of RB to remove unbound beads. The flowcell is placed in a homemade magnetic trap and a field of view with ~25-50 well-dispersed magnetic beads tethered to well-behaved, unnicked DNA molecules can readily be identified. Experiments were carried out at 27°C in RB supplemented with 0.1 mM ATP by supercoiling the DNA by 4 or 6 turns while monitoring DNA extension.

When used, RNAP roadblocks were formed on tethered constructs in RB buffer using 100-300 pM RNAP in the presence of a threefold molar excess of σ^70^ transcription factor and 0.1 mM each of ATP, UTP and GTP. Chasing RNAP roadblocks was carried out by infusing the sample with an additional 0.1 mM of CTP. Detection of transcription in the magnetic trap is well-documented and permits continuous monitoring of the presence or absence of RNAP between the mismatch and the single dam site of the DNA construct used for these experiments ^39^.

In typical incision experiments we monitored the extension of ~20 DNA molecules in parallel in a given microscope field-of-view and each molecule undergoing incision could do so typically ~5-10 times in succession before e.g. becoming stuck on the surface if supercoils accumulated over a few cycles of magnet rotation. For purposes of clarity, time-traces presented in the manuscript are aligned to the nominal end-to-end extension of DNA used given the mechanical parameters imposed.

### Surface Plasmon Resonance

60 bp duplex DNA was prepared by hybridizing 10 μM of a biotinylated [5’-bio-AGTGAGCGCAACGCAATTAATGTGAGTTAGCTCACTCATTAGGCACCCCAGGCTTTACAC-3’] and a non-biotinylated [5’-GTGTAAAGCCTGGGGTGCCTAATGAGTGAGCTAACTCACATTAATTGCGTTGCGCTCACT-3’] PAGE-purified oligonucleotide (IDT, Leuven, Belgium). The resulting duplex DNA containing Cas9 target site C was purified by gel filtration on a Superdex S200 increase column (GE Healthcare). Surface plasmon resonance (SPR) spectrometry was performed at 25°C on a Biacore T100 (GE Healthcare). A CM5 sensor chip surface was derivatized with 2000 response units (RU) of streptavidin (Invitrogen) using the amine coupling kit (GE Healthcare) and subsequently 10 RU of target site C duplex DNA was immobilized on flow cell 2. dCas9 RNA complexes containing CrRNA A or C in running buffer (25mM Hepes, 150mM KCl, and 0.05% Tween 20) were injected across the chip at 50μl/min at concentrations ranging from 0.6nM to 20nM. Absence of rebinding during dissociation was ascertained by including 100 nM of non-biotinylated duplex DNA containing target site C in the dissociation phase using the dual-injection option.

### Size exclusion chromatography

MutL WT and MutLΔ4 (1.5 mg/ml) in 25 mM Hepes-KOH [pH 7.5], 150 mM KCl, 5 mM MgCl_2_, 10% glycerol and 10 mM β-mercaptoethanol (GF buffer) were mixed with 1mM adenylylimidodiphosphate (AMPPNP) or GF buffer as control and incubated for 16 hours at 4°C. Samples (50 μl) were injected on a Superdex 200 10/300 GL column equilibrated with GF buffer and operated by ÄktaMikro (GE Healthcare) at 50 μl/min. Elution was monitored at 280 nm.

## Acknowledgements

We are grateful to Nora Goosen for the gift of BL21(DE3)pLysS,mutL::Tn10, Moara Lieuw-Hie for assistance with creation of *E. coli* strains, and Nicolaas Hermans and Alexander Fish for contributions during early stages of the project. This project has received funding from the European Community’s Seventh Framework Programme grant [223545] and Horizon2020 Marie Skłodowska Curie grant [722433]; from gravitation program CancerGenomiCs.nl from the Netherlands Organisation for Scientific Research (NWO), part of the Oncode Institute, which is partly financed by the Dutch Cancer Society, and from NWO ECHO.016.001.

## Author contributions

YM, CL, ES performed ensemble incision assays; KN, CD, AQ performed single molecule assays; CL and HW purified proteins; TKS, PF, TRS and JL designed experiments; TRS, RK and JL supervised the study; YM, TRS and JL wrote the manuscript.

## Competing Interests statement

none declared

---

**Supplemental Figure S1:**
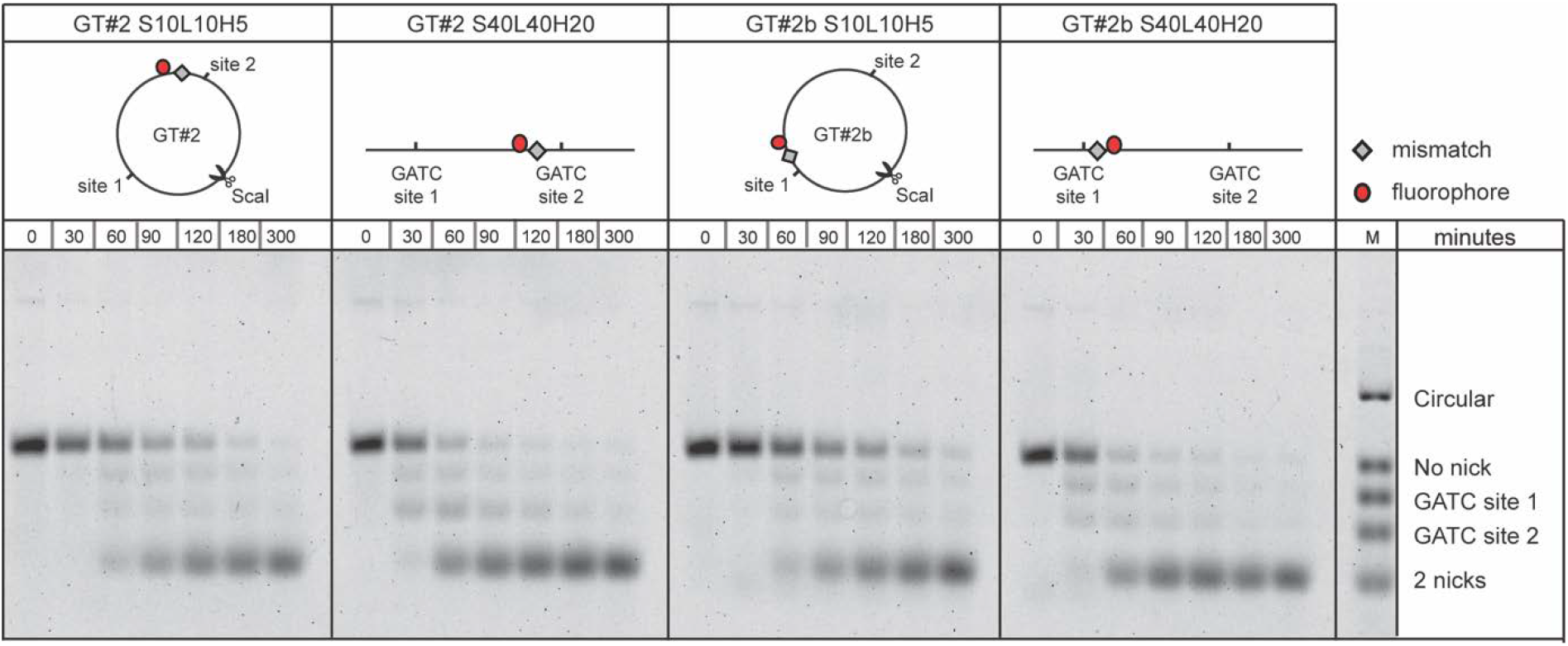
GATC site preference on linear but not on circular DNA. Time courses of GATC site incision by MutS, MutL and MutH on 0.5 nM GT#2 (2 panels on the left) and GT#2b (2 panels on the right). Reactions on circular DNA were stopped by heat inactivation followed by linearization with ScaI. Reaction products were separated using gel electrophoresis under denaturing conditions and visualized using the Alexa^647^ fluorophore.

**Supplemental Figure S2:**
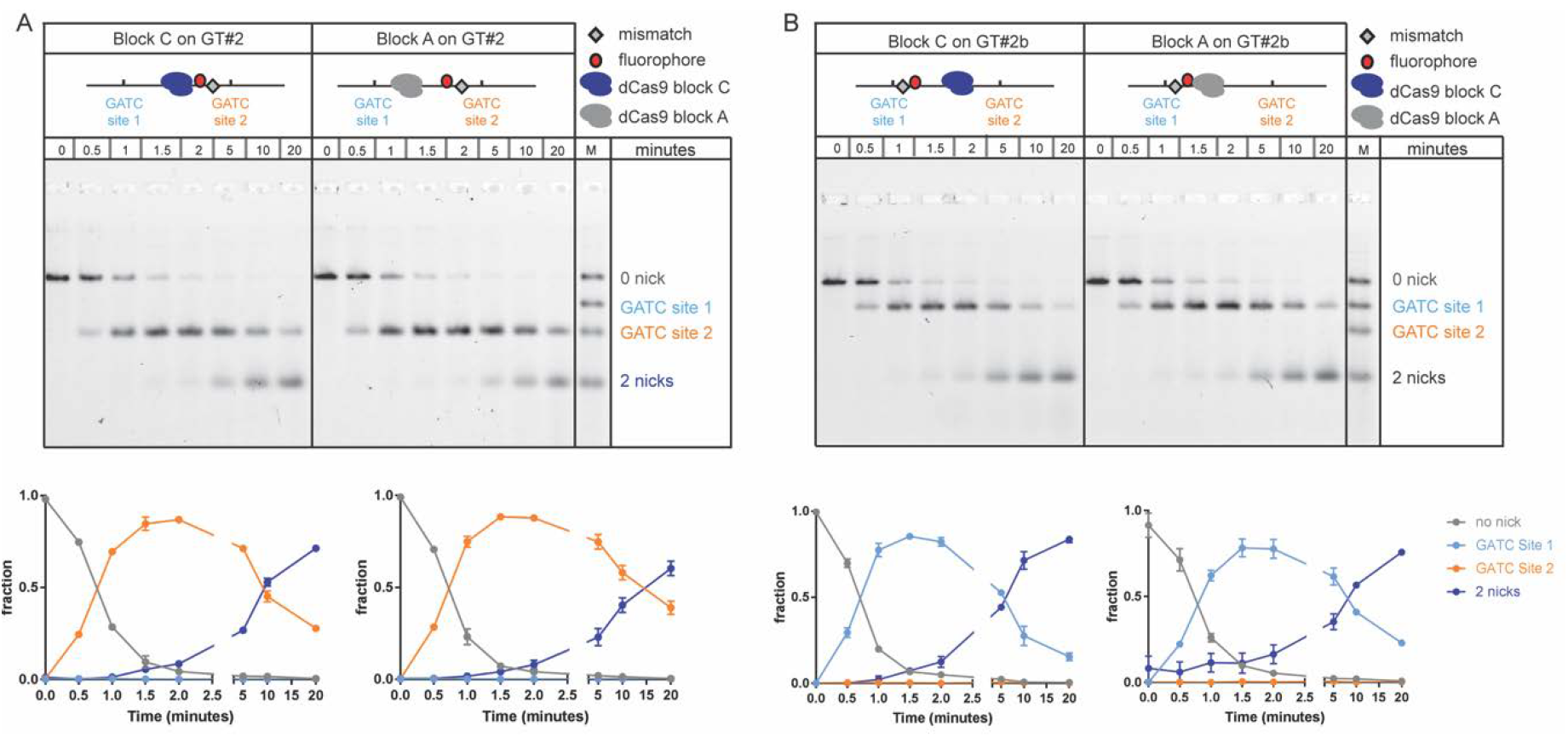
dCas9 roadblock location does not influence bypass efficiency. Time courses of GATC site incision by 50 nM MutS, 50 nM MutL and 25 nM MutH on 0.5 nM GT#2 (panel A) and GT#2b (panel B) with roadblocks at position C (blue) in the center of the substrate or position A (orange) close to GATC site 1. Reaction products were separated using gel electrophoresis under denaturing conditions and visualized using the Alexa^647^ fluorophore. Graphs show quantification of product fractions containing no nick (gray), a nick at GATC site 1 (light blue), a nick at GATC site 2 (orange) and 2 nicks (dark blue). Data points with error bars represent the mean values and range of three independent experiments.

**Supplemental Figure S3:**
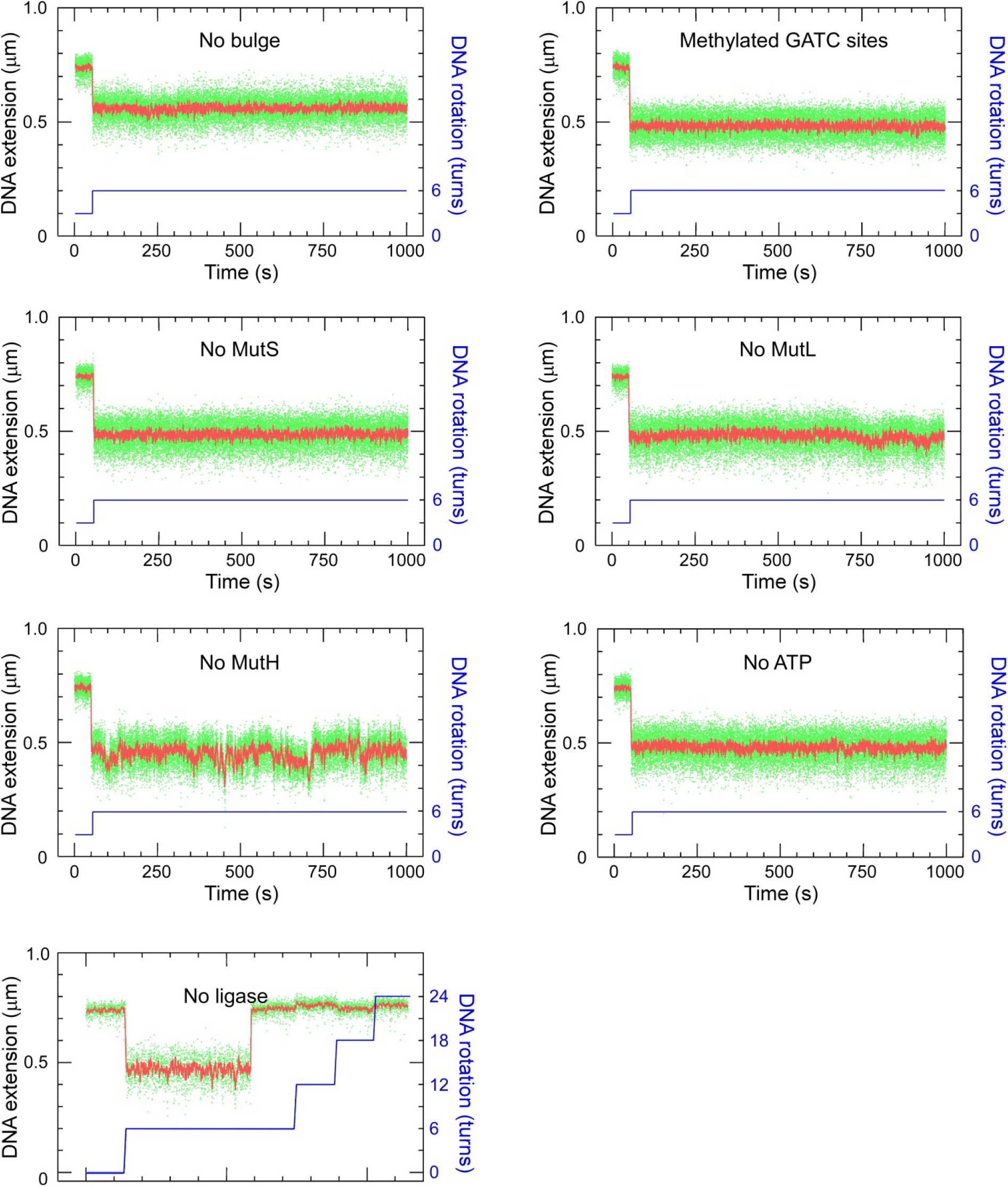
control experiments for single-molecule MMR incision assay. Time-traces obtained using supercoiled pREP4 construct show that homoduplex DNA is not a substrate for MMR (No bulge), nor are DNA molecules with methylated GATC sites (Methylated GATC sites). Removing any one of MutS, MutL or MutH (as indicated) also abolished incision, as did withholding ATP (no ATP). Finally, in the absence of T4 DNA ligase the DNA could be incised but not supercoiled again (No ligase). When present, reaction component concentrations were 2.5 nM MutS, 10 nM MutL, 10 nM MutH, and 0.1 mM ATP.

**Supplemental Figure S4:**
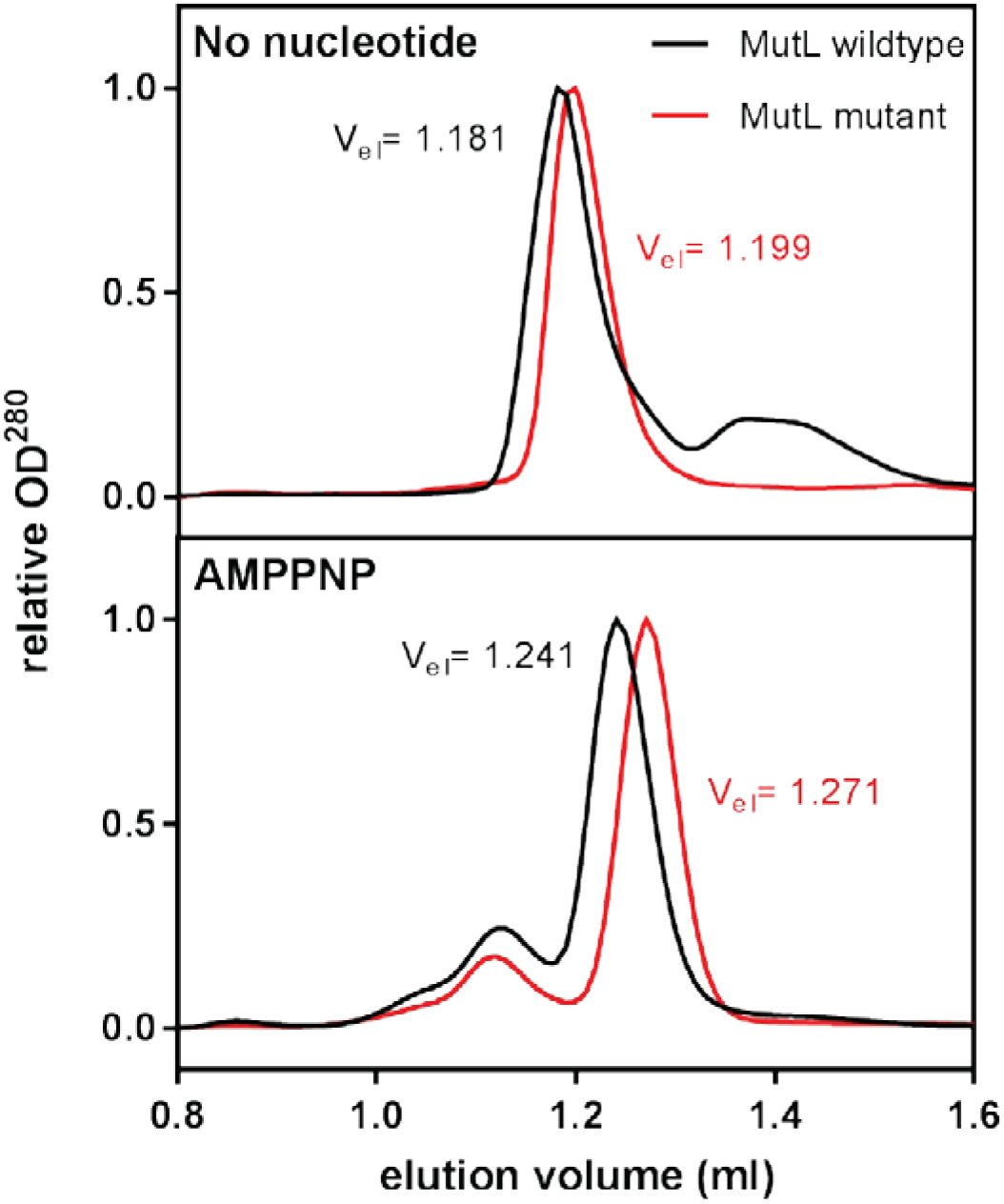
The MutL linker mutant has a smaller hydrodynamic volume than wild type MutL in the nucleotide-free and -bound states. Upper panel: Normalized size exclusion chromatography data of wild-type MutL (black) and MutLΔ4 (red) in open form. The increase in elution volume (V_el_) indicates a smaller hydrodynamic volume for MutLΔ4 compared to wild type MutL. Lower panel: Wild-type and MutLΔ4 MutL in the closed state, induced by the presence of nucleotide AMPPNP show a similar difference in hydrodynamic volume.

